# Motility-gradient induced elongation of the vertebrate embryo

**DOI:** 10.1101/187443

**Authors:** Ido Regev, Karine Guevorkian, Olivier Pourquie, L Mahadevan

## Abstract

The body of vertebrate embryos forms by posterior elongation from a terminal growth zone called the Tail Bud (TB). The TB produces highly motile cells forming the presomitic mesoderm (PSM), a tissue playing an important role in elongation movements. PSM cells establish an anterior-posterior cell motility gradient which parallels the degradation of a specific cellular signal (Fgf8) known to be implicated in cell motility. Here, we combine electroporation of fluorescent reporters in the PSM to time-lapse imaging in the chicken embryo to quantify cell diffusive movements along the motility gradient. We show that simple microscopic and macroscopic mechano-chemical models for tissue extension that couple Fgf activity, cell motility and tissue rheology at both the cellular and continuum levels suffice to capture the speed and extent of elongation. These observations explain how the continuous addition of cells that exhibit a gradual reduction in motility combined with lateral confinement can be converted into an oriented movement that drives body elongation. The results of the models compare well with our experimental results, with implications for other elongation processes in the embryo.

---

Most vertebrate species exhibit an elongated body axis. This characteristic pattern is established during embryogenesis as the tissues progressively form in an anterior to posterior direction. Microsurgical ablation of the posterior PSM (which contains the precursors of skeletal muscles and axial skeleton), severely reduces posterior elongation movements, indicating that this tissue plays a major role in the control of posterior elongation of the embryonic axis (Fig 1(A,E)). Analysis of cell motility in the chicken embryo PSM [1] shows that there is an antero-posterior gradient in the activity of cells. However, locally, the motility inside the PSM of chicken and zebrafish embryos is manifested by random, undirected, Brownian-like cellular motion [1, 13]. These random, diffusive movements contrast with the oriented cell intercalation movements controlling elongation of the anterior parts of the embryo [13]. Furthermore, inside the PSM, this gradient is downstream of a gradient of the secreted growth factor Fgf8, as shown schematically in Fig. 1(A). Fgf8 is known to play an important role in cell motility [6]; indeed, increasing Fgf8 concentration in the PSM causes cell motility to increase without any orientation preference [1], reducing the speed of elongation. While earlier qualitative models focused on the role of cell addition at the TB[1], how this motility gradient generates a driving force in an elongating tissue remains unstudied.

**Fig. 1.**
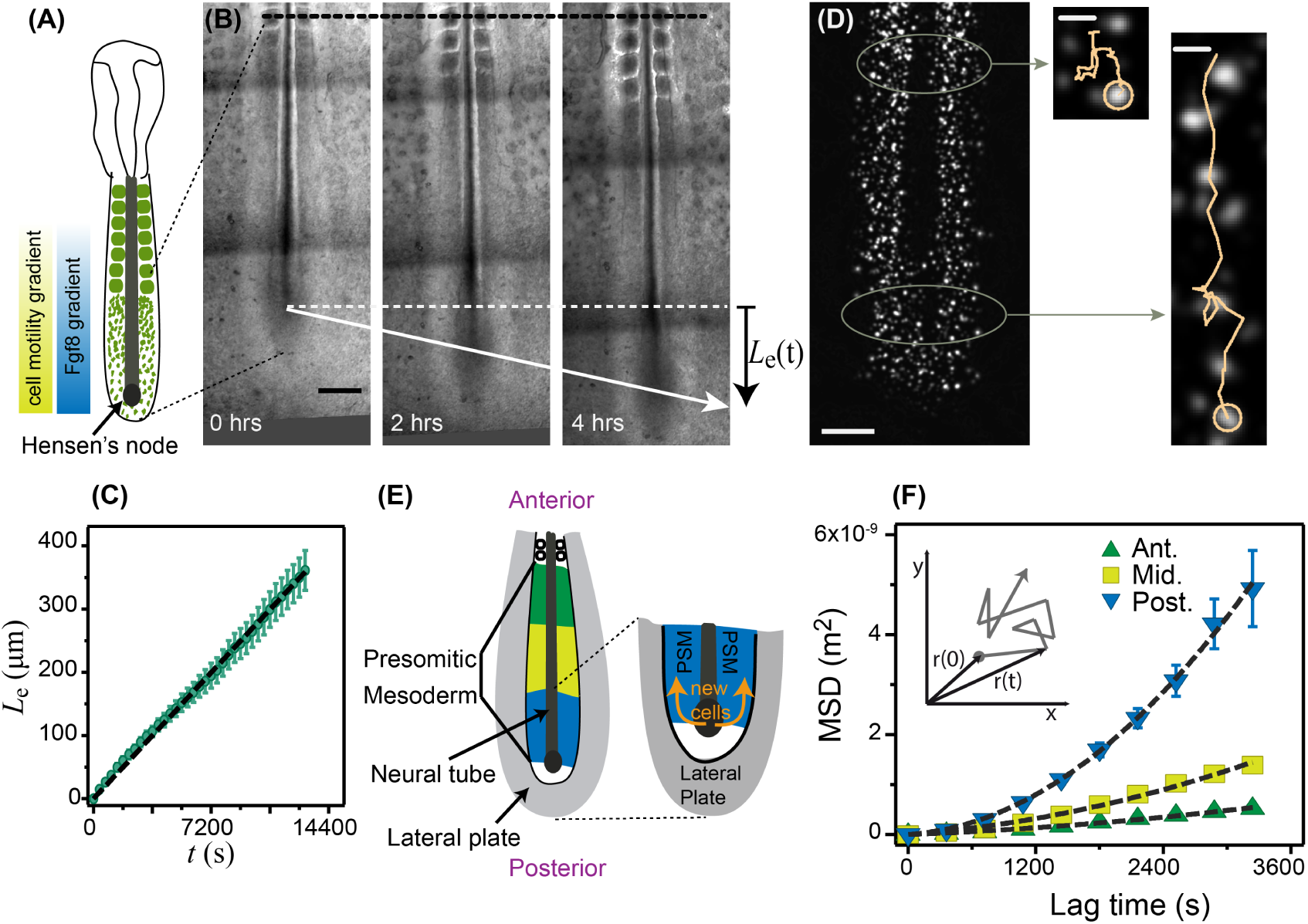
Axis elongation and cell diffusion in chicken embryo. (A) Schematic of an embryo at stage 10 of Hamburger-Hamilton. Cell motility decreases from posterior to anterior, in correlation with a decrease in the Fgf8 concentration. A gradient of cell density opposite to the motility gradient is shown in the schematic embryo (in green) (B) Time series of an elongating embryo. The black dotted line shows the reference point for tracking the posterior elongation. *L*_*e*_*(t)* is the distance that the Hensen’s node advances over time. Scale bars represent 200//m. (C) Elongation of the PSM, as a function of time. The slope gives the average elongation rate *V =* (2.8 ± 0.3) × 10^-2^ μs (n=5, mean ± SEM). (D) Electroporated cells inside the PSM. Anterior cells advance much less than the posterior cells for the same duration of time (here 4 hours). Scale bars are 200μm for whole PSM and 25/im for zoomed tracks. (E) A schematic of the PSM showing the three regions considered for MSD analysis shown in (F), as well as the depiction of the neighbouring tissues. New stem cells are generated by division of the progenitor cells inside the TB, and move into the PSM. The movement of cells in the PSM is limited by the neural tube medially, the somites anteriorly and the lateral plate laterally. (F) Average MSD for the anterior, middle and posterior PSM. Dashed lines are adjustments by Eq.1. Inset shows a sketch of random motion.

The observation that locally, cell motility is undirected needs to be reconciled with the emergence of oriented cellular motion and posteriorly-oriented cellular motion leading to body elongation. A potential mechanism for the observed directional cell velocity is thought to be the inhomogeneous mechanical pressure associated with cellular motility that leads to the rectification of cell motion and to forces that cause the tissue to elongate. Because the expression of Fgf8 is highest at the posterior PSM, and Fgf8 expression decreases away from it, this leads naturally to a reduction in motility anteriorly until the effects of adhesion cause the cells to eventually condense into somites. This hypothesis is consistent with observations of outgrowths in other morphogenetic situations such as in the limb bud [8, 22]. As new cells enter the PSM they are exposed to a high concentration of Fgf8 and become highly motile, but do not move in an oriented manner. When combined with the confinement due to the presence of relatively immobile and stiff lateral tissues [2], this yields an effective pressure that causes the body to extend. As Fgf8 degrades over time, anteriorly positioned cells move less, before eventually coming to a rest as they aggregate into epithelial somites. To better understand these processes, here we use quantitative experimental observations to measure the effective diffusivity of cells as well as their elongational speed as a function of their location relative to the last formed somite. Our observations suggest a minimal microscopic cellular description of a zone of proliferating cells with high motility that we use to develop a quantitative cellular model, and an equivalent macroscopic continuum description of body elongation based on the cellular model and experimental observations. Together, these complementary approaches yield simple expressions for the speed and scale of body elongation consistent with our experimental measurements, with implications for our understanding of outgrowth morphogenesis in other settings such as limb and gut formation.

## I. EXPERIMENTAL OBSERVATIONS

Our experiments were carried out with chicken embryos at HH stages 10-11 [10], corresponding to the period when the elongation of the embryo is most substantial [7]. A time-series of an elongating PSM is shown in Fig. 1(B) (Supp. Movie 1). To measure the elongation rate we register the movement of the embryo with respect to the last formed somite at the beginning of the experiment as depicted by the black dotted line in Fig. 1(B), and track the advancement of the Hensen’s node, *L*_*e*_ *(t)* as a function of time. In Fig. 1(C), we see that *L*_*e*_ *(t)* increases linearly with time, with a mean elongation rate *V* = (2.8 ± 0.3) × 10-^2^ μs (averaged from five embryos).

To evaluate the role of cell motility on overall body elongation, we now turn to study the movement of cells by electroporating the PSM cells with fluorescent reporters specifically labeling cell nuclei (H2B-GFP) [11], as shown in Fig. 1(D) (Supp. Movie 1). As a first approximation, the motion of the cells can be considered two dimensional in the antero-posterior and medio-lateral directions because the relative ventro-dorsal depth of the PSM is small. In the reference frame of the last fixed anterior somite, during a fixed acquisition time of 4 hours, posterior cell trajectories show a larger net displacement than the anterior ones, consistent with prior experiments [1]. To quantify the variations of cell motility along the body axis, we first divide the PSM into three regions: anterior, middle and posterior as shown schematically in Fig. 1(E), and obtain the mean square displacement (MSD) of the cells given by 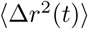, where 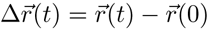 defines the distance that the cell travels in a lag time t, as shown in the inset of Fig. 1(F) [19, 23]. Decomposing the motion into a random diffusive term and an oriented drift term[17], we write the MSD as:

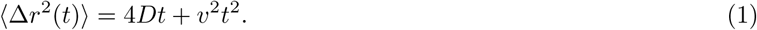

where *D* is the effective population diffusivity and *v* is the local population drift velocity [19]. This model for cell dynamics is in accordance with previous findings for chicken [1] and zebrafish [13] PSM elongation. Figure 1(F) shows the mean MSD curves of the three regions (anterior, middle, posterior) defined in Fig. 1(E) and the fit to Eq. 1 (black dashed curve). From the fits we obtain *D*_*Post*_ = (3.5 ± 0.7) × 10^-2^ μ^2^, *D*_*Mid*_ = (2.1 ± 0.4) × 10^-2^ μm^2^/s, *D_Ant_* = (1-4 =b 0.3) × 10^-2^ μm^2^ and *v*_*Post*_ = (2.0 db 0.2) × 10^-2^ μs, *V_Mid_* = (1-1 ± 0.1) × 10^-2^ μs (mean db SEM), *V*_*Ant*_ = (0.6 ±0.1) × 10^-2^ μs. These estimates confirm the presence of a motility gradient of cells along the AP axis.

All together, our observations quantify how the cells move in space and time along the PSM. When new cells are added close to the tailbud, they are highly motile but move in random directions. As they move further away from the zone of proliferation in the TB they gradually slow down and stop moving. We now turn to quantify the gradient in motility coupled with the confinement provided by the (more rigid) lateral tissues and show that this suffices to direct cell motility and explain body elongation as a chemo-mechanical process.

### II. MICROSCOPIC CELLULAR MODEL

We start with a simple cellular model built to mimic the experimental observations in a quasi-one dimensional setting. Our simplifying assumptions, which are used in both cellular and continuum models, are that the width of the PSM is constant, which is known to be approximately true for the region where cells are motile[2], and that the cells cannot escape from the PSM owing to the constraints imposed by the somites anteriorly, the lateral plate laterally and the neural plate medially. The assumption of quasi-one dimensional dynamics is manifested in the simulations by using periodic boundary conditions in the direction perpendicular to the direction of elongation instead of rigid-walls and in the fact that the density only depends on the elongation direction, in the continuum model. In the discrete model, we model individual cells as soft, elastic, disks which move randomly in a manner analogous to a Brownian particle, recognizing that the cause of random movement is not related to the temperature of the environment but instead corresponds to the active but random motility of the cell [3, 14]. The equation of motion for a cell with coordinates *r_i_* is:

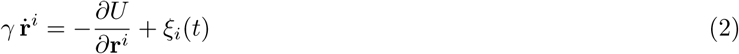

where *γ* is the viscous friction and we have assumed that inertial effects are negligible, so that we can consider overdamped motion. The viscous friction is a result of the interaction of cells with the extra-cellular matrix (ECM)). We assume that there is a short range repulsive interaction between cells of size *a* to guarantee that two cells cannot occupy the same position and that the potential is harmonic, being given by:

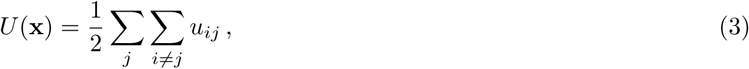

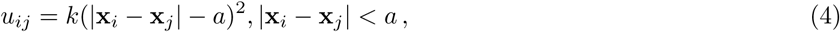

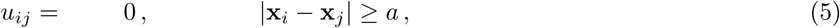

Note that in our simulations the cell-cell excluded volume interactions are conservative and thus collisions are elastic but overdamped. The random force *ξ*(*t*) is assumed to be zero-mean and normally distributed with Gaussian statistics so that:

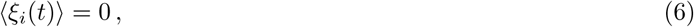

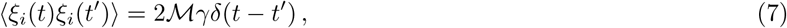

where *M* is the single-cell activity/motility. In the following, we will also use a result from statistical physics [12], that the microscopic diffusivity of a (Brownian) cell is related to the activity by the relation 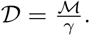 Using the equations for the motion of cells, we simulate their dynamics by ensuring that each cell that is injected at the PSM boundary starts with the same initial activity; at each time step, we turn off the activity (and thus decrease the fraction of motile cells) with a probability which increases exponentially with time:

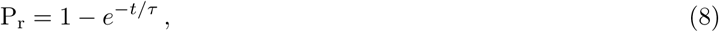

where *τ* is the slowest time scale associated with kinetics of degradation of Fgf8 that initiates once the cell enters the tailbud. In the following we will assume that for a given simulation, Fgf8 levels and thus *τ*, are constant, which is approximately true in the experimental setup for the observed times of 4Hrs. We consider the dynamics of these cells in a quasi-one-dimensional situation wherein the cells are confined between two rigid lateral walls (we assume that the lateral plate is relatively stiff [2]), and further limited by the anterior region where the somite are stationary and cells are non-motile. Since we are interested in the quasi-one dimensional case we use periodic boundary conditions instead of rigid walls for the lateral plate (in both periodic and rigid boundary conditions cells cannot escape through the boundaries). At the posterior end corresponding to the tailbud, we assume that cells can move as the body elongates owing to a constant rate at which they are added in the space that is not occupied by other cells (Fig 2(A,B)). The motivation for this assumption is the fact that cells are generated when the progenitor cells in the TB divide, enter the PSM and become motile (Fig 2(A)). The movement of the cells from the PSM to the TB region is limited by the available space in the PSM (the TB itself is not modeled in our quasi one-dimensional model). Therefore, also in our model, if there is no free space, no cells are added, so that the rate of adding cells is limited by the motion of the PSM and the maximal cell packing density *ρ*_0_. This implies that cells inside the TB divide in response to cell migration from the TB to the PSM. While there are no direct evidence for this in embryo elongation, similar behavior was recently observed in skin tissue [16]. The tailbud boundary is modeled as a wall of immobile cells attached to their neighbors by elastic springs with spring constants *k*_*chiain*_ = 2k, and allowed to move posteriorly due to the mechanical pressure exerted by the motile cells anterior to it. Our simulations show that after an initial transient, the motion of the wall reaches a steady-state where cells added at a constant rate cause the wall to move at a constant velocity (Supp. Movie 2). The fraction of motile cells is much larger near the moving wall, and as one moves anteriorly, this density falls off quickly due to the decrease in the activity of Fgf8. Changing Fgf8 activity by changing τ and thus varying the probability with which each individual cells stops moving (and thus the total fraction of moving cells), changes the velocity and motility profiles (see figure 2(C,D)).

**FIG. 2.**
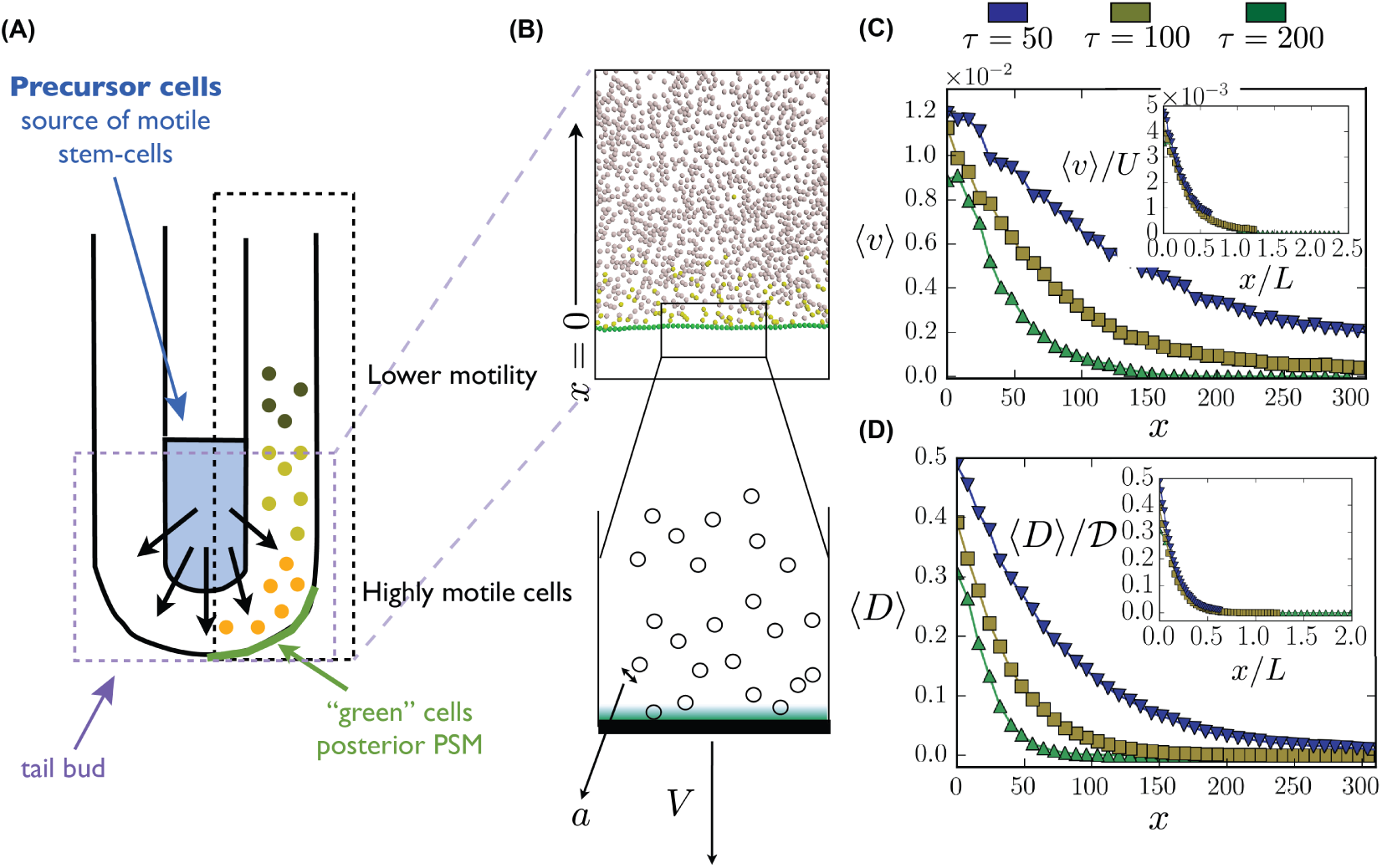
Microscopic cell-based simulation: (A) Schematic view of the Presomitic mesoderm used as a basis for a cellular simulation. (B) Yellow and grey spheres represent motile and immotile cells respectively. The green spheres form a connected wall which represents the tailbud and can move in response to pressure applied by the motile cells. The wall velocity is *V* and the size of the cell is *a*. (C,D) Velocity and motility profiles for different τ values are calculated as in the experiments, by fitting to Eq. 1. *Inset:* scaling *x* by L, *(v)* by *U* and *(D)* by *T)* shows that the curves collapse onto each other demonstrating that *L* is the relevant length-scale over which cells are motile. Here we express *v, T>* and *x* in terms of dimensionless units as described in our simulations (see Materials and Methods). Here we used *D* = 2.5, *k* = 100, *α* = 1 and γ 50 in the simulation units.

To understand these numerical results qualitatively, we note that a new cell can be added at the (moving) boundary only when there is a gap of order of the size of one cell *a* (see Fig. 2(B)). This allows us to draw an analogy to the Brownian ratchet problem which was introduced in the context of polymerization [18]. In the PSM region where the internal tissue resistance to cell motion dominates any external resistance, as in our simulations, the rate of elongation is limited by the waiting time for a gap to open that allows for the addition of a cell of size *a* i.e. 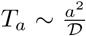. Here *D* is the activity (diffusivity) of a single cell, which is different than the population-averaged diffusivity of the cells *D* (consistent with Eq. 1). In this limit of diffusion-limited elongation (*T*_*a*_ ≪ t), the speed of elongation scales as *a*/*T*_*a*_, i.e.

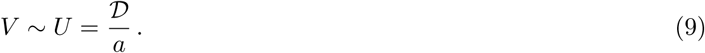

Similarly, the length scale over which the fraction of motile cells falls off exponentially is

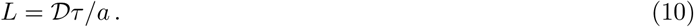

We note that the other limit, i.e. *T* ≪ *T*_a_ is tantamount to adding cells that are not active, a situation that will lead to a jammed state, and thus not relevant here. In Fig. 2(C,D) and insets, we show, by rescaling the velocity, cell diffusivity and distance, that the observed dependence of the speed of elongation and the effective diffusivity as a function of location from the wall is consistent with our simple scaling arguments. However, as can be noticed in the inset of Fig. 2 C, although the wall velocity in our simulations *V* ∼ *D/a*, it is much larger than the cell drift velocity *(v)*. This is because we used large values of the cell diffusivity to speed up our computations.

## III. MACROSCOPIC CONTINUUM THEORY

To complement the statistical model for the population dynamics of cell motility considered in the previous section, we now turn to an effective macroscopic continuum theory that links the density of motile cells *ρ*(*x, t*) and the velocity field *ρ*(*x, t*) of motile cells as a function of location *x* in a fixed lab frame, similar to that used to describe the dynamics of active fluids [15]. A hydrodynamic description of the diffusion, advection and degradation of motile cells can then be written in terms of the equations for mass and momentum balance as:

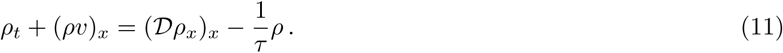

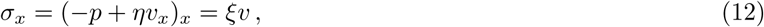

This first equation describes mass balance for the density of motile cells that moves, diffuses and degrades, while the second one characterizes how an active pressure generated by the motile cells causes them to exert forces on each other and thence the tailbud. Here, we have used the simplest linear relation linking the active pressure to the density of motile cells *ρ* ∼ *αρ*, consistent with the microscopic model with *a* ∼ *M*. (we have also considered other relations of the form *p* ∼ *αρ*^*q*^ but found that *q =* 1 provides the best fit to the data and is consistent with simulations of active Brownian particles[14]), and assumed that *η* is the viscosity of the fluid of motile cells and *ξ* is the viscous friction associated with motion of the elongating tissue relative to the surrounding tissue (endoderm, ectoderm, neural tube and lateral plate), and finally have neglected any inertial contributions.

To complete the formulation of the problem, we need to specify some boundary conditions for the free-boundary problem. The position of the tailbud where new cells enter the domain is assumed to be *s(t)* so that the domain of interest is *x* ∈ [*s*(*t*), ∞). Assuming that motile cells enter the domain at a rate proportional to the difference between their local density (at the boundary all the cells are motile) and the maximum cell density *ρ*_0_, we write the cell flux as *R*(*ρ*_0_ - *ρ*(*s*)). This flux must be balanced by diffusion and advection of cells from the boundary so that

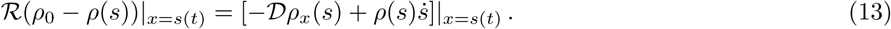

which is reminiscent of a generalized Stefan-like condition in moving boundary problems in solidification. We also have to satisfy force balance at the moving boundary so that

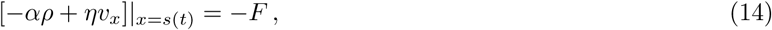

where *F* is the resisting pressure exerted by the tissue ahead of the tailbud, and *v*(*s*) = *ṡ* (in the cellular simulation, this force is a result of the dynamic friction between the wall cells and the substrate and thus depends on the velocity, here we assume a more general force). Far from the tailbud, we assume that due to degradation of Fgf8, the density and velocity of motile cells vanishes so that:

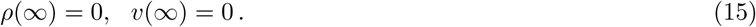

Together, (11-12) along with the above boundary conditions determine the spatio-temporal evolution of the density and velocity fields in the elongating embryo as well as the speed of elongation of the embryo itself, which will be found by solving the equations self-consistently.

In a steady-state, relative to a co-moving frame with an origin attached to the elongating tip, the PSM boundary is stationary, while the anterior moves at an unknown speed -*V*. Then we can write Eq. 11-12 at the moving frame, in which it reduces to:

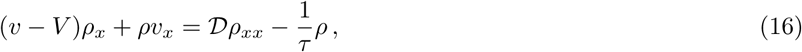

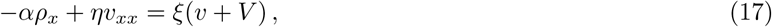

together with the boundary conditions:

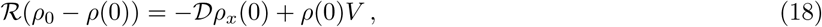

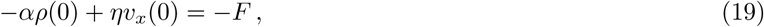

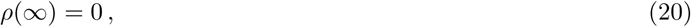

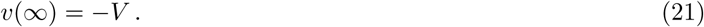

To understand the dependence of the solution of Eq. (16-21) on the problem parameters we rewrote the equations in a form that depends on only five dimensionless parameters that are a combination of the parameters of the problem:

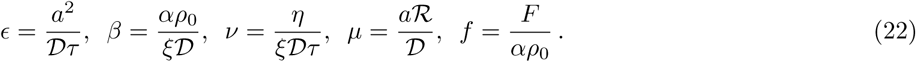

Some of these parameters were known and some were obtained by fitting as will be explained below. The first parameter, *ε* = *a*/*L* ≪ 1 links the microscopic picture characterized by cell size to the macroscopic picture characterized by the elongation length scale *L* = *D*_*τ*_/a. From the experiments, we find that the measured diffusivity at the tailbud is *D* ≈ 0.1 μm^2^/s, while the typical degradation time-scale was found to be *τ* ≈ 2 × 10^4^ s. Assuming that a typical cell size of *a* ≈ 10μm, we find that *ε* ∼ 0.05, consistent with our approximation that body elongation is dominated by the diffusively limited process of adding cells. Using the results of previous measurements of tissue rheology using a micropipette aspiration technique [9], we estimate the viscosity of the PSM to be of the order of *η* ∼ 10^4^ Pa • s. We solve the coupled set of nonlinear equations using the MATLAB© procedure BVP4C to find the velocity *V*, the density of motile cells *ρ*(*x*) and velocity *v*(*x*) self-consistently. We have found that the best agreement with the experimental data was obtained for the dimensionless parameters values *β* ≈ 5000, *v* ≈ 2000, *μ* « 3 and *f* ≈ 0.001.

In Fig.3 B,C we show the experimentally determined moving average of *v* and *D* = *D_ρ_*/*ρ*_0_ as a function of position relative to Hensen’s node (see Materials and Methods), showing a continuous decrease of both quantities from posterior to anterior of the PSM and see that these results for the elongation velocity and effective motility profiles compare well with our continuum model (given the choice of our dimensionless parameters). Our scaled results in the insets of Fig. 2C,D show that we can qualitatively capture the dynamics of body elongation driven by gradients in cell activity. Our calculated profiles for the velocity *v*(*x*) and diffusivity *D*(*x*) = *T*>p(*x*)*ρ*_0_ show a decaying form suggestive of a simple exponential law. However, careful consideration of the different terms in Eq. [16-21] shows that while the diffusive term associated with the density is small, the nonlinearities are not negligible and so our solution for the density of motile cells (and the velocity) is not quite exponential.

**Fig. 3.**
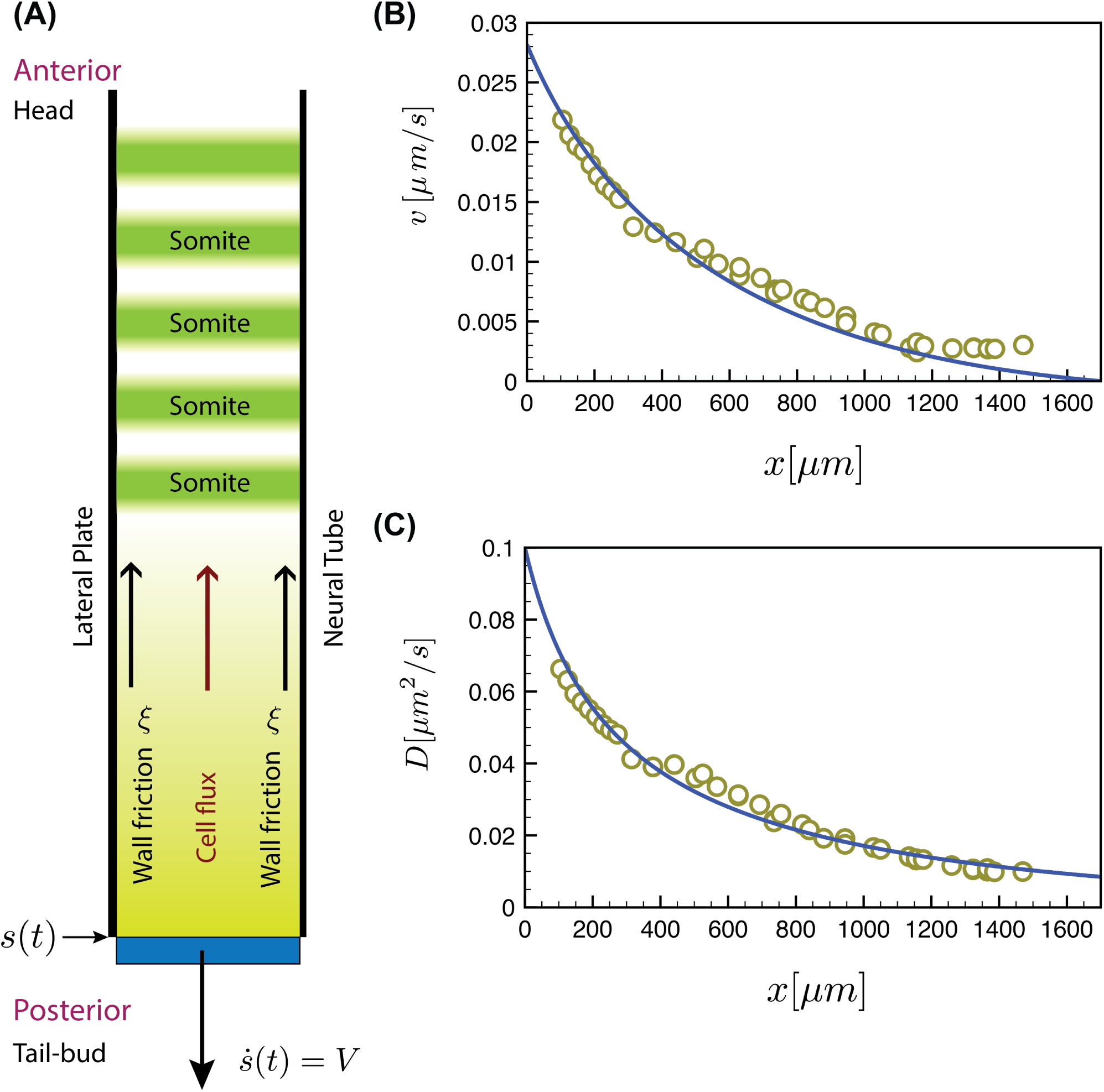
Macroscopic continuum model (A) Schematic showing half of the PSM in the neighborhood of the tailbud whose position is s(t).(B C) Experimentally measured velocity *v*(*x*) and diffusivity *D*(*x*) profiles (calculated using Eq. [1]) as a function of distance from the tailbud compares well with the results of our continuum theory obtained by solving Eq. 11-12 for *v*(*x*) and *D*(*x*) = *D*_ρ_(*x*)/ρ_0_ (continuous lines).

From solving the equations self-consistently we also find that *V* ≈ 0.03μl;m/s. We note that *f* ≪ 1 0ndicates that *F* ≪ *αρ*_0_ and thus the pressure resisting the tailbud motion is negligible compared to the pressure exerted by cell motility. This also means that the elongation process depends quantitatively only on three dimensionless parameters; the scaled activity *β* the scaled addition rate *μ* and the scaled viscosity *v*. When combined with our choices for the dimensionless parameters, this allows us to estimate the friction coefficient *ξ* ≈ 0.7 Pa × s/m^2^, the active stress *αρ*_0_ ≈ 1.323 Pa and the force resisting tailbud elongation *F* ≈ 1.45 × 10-^3^ Pa. These values can serve as a reference for future measurements on amniote embryos [21, 24].

## IV. DISCUSSION

In vertebrate embryos posterior structures are formed sequentially by a combination of cell proliferation and cell motility that together leads to body elongation. While it has long been observed that the elongation process involves the posterior displacement of the tailbud with respect to the head, the physical mechanism that allows for this elongation process to take place has not been clear. Here we have quantified this process in the context of body elongation and shown that it occurs as a result of two effects: the addition of motile cells at a boundary (the TB) that leads to forces generated by the rectification of random cell diffusivity by confinement, yielding a characteristic *velocity scale* and *length scale*. Given that other embryonic outgrowths such as the vertebrate limb bud also exhibits graded diffusive behavior of its cells downstream of Fgf8 signaling [8], the mechanism we propose for elongation might be more widely applicable in a variety of different contexts in vertebrate morphogenesis.

## V. MATERIALS AND METHODS

### A. Chicken embryo preparation and electroporation

Fertilized chicken eggs were obtained from a commercial provider (Les Couvoirs de l’Est, Willgottheim, France) and incubated at 37°C in a humidified incubator. After 24 hours, stage 4-5 HH (Hamburger-Hamilton [10]) embryos were mounted on filter paper and transferred ventral side up to 35 mm agar/albumen petri dishes for injection [4]. The electroporation of the PSM was performed using H2B-mCherry or H2B-Venus nuclear markers as described previously [1]. Electroporated embryos were returned to incubator and left to grow to 10-11 HH stage before imaging.

### B. Time-lapse imaging and track analysis

The imaging procedure used here is similar to previously described procedure [5, 20]. Briefly, the embryos were transferred to custom made six-well observation chambers containing agar/albumin gel, and positioned ventral side up. The time-lapse imaging was performed at 37°C using a motorized upright microscope (Leica DMR, Leica Microsystems) with a 10x objective (N.A. 0.3) and a CCD digital camera (QImaging Retiga 1300i) at 10 frame/hr rate.

At each time-point, bright-field and fluorescent images of the embryo were taken at 3 various fields to cover the total length of the PSM. Post acquisition processing were performed on images as described previously [5, 20] to obtain a 2D time-series. Cell tracking and trajectory analysis was performed on fluorescent images using custom made Matlab (MathWorks) routines. For each cell trajectory, the MSD is calculated and adjusted with Eq.1 to obtain *D* and *v*. Each data point in Fig.3 is obtained using the Smooth function in Matlab.

### C. Simulations

The simulations were performed by discretizing Eq. 2 using the Euler-Maruyama method with a Langevin term and integrating in time. As usual for this kind of simulations, we use reduced, dimensionless units in which all energies are given in terms of a typical energy *w*: E^*^ = *E/w*, all lengths in terms of the typical cell size a, *r*^*^ = *r/a* and masses in terms of a mass unit *M*, *m*^*^ = *m*/*M*. Using these fundamental units we can also reduce the temperature (or motility) *T*^*^ = *k*_B_*T*/*w*, time 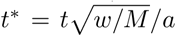 fraction *ρ*^*^ = *Na*^3^/*V* (*V* is the volume) and any other physical quantity of interest, where *E*^*^, *r*^*^, *m*^*^, *T*^*^ and *ρ*^*^ are all dimensionless.

## ACKNOWLEDGMENTS

This work was partially supported by the French Agence Nationale pour la Recherche under grants No. ANR-14-CE32-0009-01 to K.G., Human Frontier Science Program RGP0051/2012 to O.P., and grants and fellowships from the Schlumberger Foundation and the MacArthur Foundation to L.M.

